# A Suite of Stains: Characterization of four fluorophores as complementary tools for visualizing neutral lipids in an extremophilic green alga

**DOI:** 10.1101/2025.08.18.670973

**Authors:** Pomona Osmers, Eliza-Jayne Y. Boisvert, Christopher N. Boddy, Deryn E. Fogg, Marina Cvetkovska

## Abstract

Understanding lipid metabolism in algae is critical to advancing our knowledge on fundamental algal physiology and for harnessing these organisms as platforms for the sustainable production of high-energy lipids. BODIPY is the most prevalently used fluorescent dye for the visualization of lipid droplets (LDs) in algae; however, its limitations warrant exploration of alternatives. Here we evaluate and compare four lipophilic fluorophores (BODIPY, DAF, Cou, DPAS) for their effectiveness in staining LDs in the extremophilic green alga *Chlamydomonas priscui*. We assess each dye’s photophysical properties, synthetic accessibility, LD specificity, cellular toxicity, and suitability for microscopy and flow cytometry. All four dyes successfully stain LDs, but their performance diverges under different experimental conditions. BODIPY permits long-term incubation allowing quantification in time-course studies but exhibits poor LD specificity and high susceptibility to photobleaching. DAF enables polarity-sensitive staining but is highly toxic on prolonged exposure or during cellular stress. Cou and DPAS yield strong LD-specific signals with low cytotoxicity, making them ideal for studies involving environmental stress. However, DPAS requires room-temperature incubation, pointing toward greater potential utility for non-extremophilic algae. These results expand the toolbox for lipid biotechnology research in extremophiles and underscore the importance of tailoring dye selection and experimental conditions to algal physiology.

## Introduction

As we are nearing the end of the petrochemical era, sustainable access to renewable resources is urgently needed. Plant- and algal-sourced triacylglycerols (TAGs) have long been valued as sources of food and biofuels and, more recently, as substrates in the oleochemical and pharmaceutical industries (Biermann et al., 2021; Long et al., 2021; Lu et al., 2011; Montero de Espinosa and Meier, 2011). Aquatic algae have many advantages over land plants for TAG production. They exhibit rapid growth, superior photosynthetic efficiencies, can be cultured on non-arable land using saline and/or wastewater, and have the potential to produce significantly more lipids per acre than land plants (Fabris et al., 2020; Wang et al., 2024). Research on TAG biosynthesis and turnover in algae is hence of utmost importance. The green alga *Chlamydomonas reinhardtii* has emerged as one of the leading models for understanding algal TAG accumulation (Merchant et al., 2012; Scranton et al., 2015).

In green algae, TAGs are synthesized via a complex biosynthetic pathway and stored in specialized organelles termed lipid droplets (LDs) (Li-Beisson et al., 2021; Li-Beisson et al., 2023). LDs are highly dynamic and have a myriad of functions. In addition to serving as cellular storage depots for high-energy neutral lipids, these organelles prevent lipotoxicity arising from the accumulation of free fatty acids, participate in signal transduction, and maintain membrane homeostasis (Ischebeck et al., 2020). Typically, TAG-rich LDs form in algal cultures that are not actively growing and have a surplus of carbon and energy: in cells in the stationary phase, under conditions of nutrient deprivation (Davey et al., 2014; Juergens et al., 2016; Moellering and Benning, 2010; Siaut et al., 2011; Urzica et al., 2013), and in response to environmental stress (Goold et al., 2016; Hemme et al., 2014; Légeret et al., 2016). In turn, LDs are degraded when algal cells enter periods of rapid cellular division and growth fueled by the energy-rich acyl chains (Jüppner et al., 2017; Lee et al., 2020; Tsai et al., 2015). While this dynamic is crucial for maintaining cellular homeostasis, it poses a key challenge for biotechnological application. Stressful conditions promote lipid production but reduce growth, limiting the potential for continuous and efficient TAG production.

Cold-water extremophilic algae from polar environments have the capacity to fill this gap, and have drawn recent attention as novel biotechnology platforms (Malavasi et al., 2020; Morales-Sánchez et al., 2020b; Varshney et al., 2015). Green algae are some of the best understood cold-water eukaryotes known to thrive under conditions untenable for temperate species, including permanently low temperature, low light, and high salinity. These conditions promote the biosynthesis of lipids with a high polyunsaturated fatty acid (PUFA) content, as well as carbohydrates and antioxidants, as natural adaptive mechanisms that enable robust growth under extreme conditions without sacrificing biomass (Cvetkovska et al., 2017; Hulatt et al., 2017; Hüner et al., 2022). While the lipid content of polar *Chlamydomonas* has been previously examined by GC-MS (Ahn et al., 2015; Jung et al., 2016; Kim et al., 2016; Morales-Sánchez et al., 2020a; Morales-Sánchez et al., 2020b; Zhang et al., 2011), there are currently no insights on LD dynamics and lipid turnover in these species.

The use of fluorescent, lipid-specific dyes enables rapid detection of neutral lipids and visual investigation of the size, number, localization, and dynamic behavior of LDs within algal cells. The commercially available fluorophore boron dipyrromethene (BODIPY; Ex/Em = 505/515 nm) is one of the most widely used fluorescent dyes in algal studies (reviewed in Patel et al., 2019; Rumin et al., 2015). BODIPY has several advantages over earlier-generation lipid stains such as Nile Red (Alemán-Nava et al., 2016), including improved LD specificity, photostability, and rapid staining. BODIPY also effectively penetrates the algal cell walls in model algae, although low stain permeation rates are reported in species with thick and/or rigid cell walls (Brennan et al., 2012; Südfeld et al., 2021). Other important features include broad excitation and emission maxima in the blue and green regions of the visible spectrum (450-505 nm and 515-550 nm, respectively), enabling excitation with widely accessible blue lasers, and resulting in emission that is spectrally distinguishable from the red autofluorescence of the algal chloroplast (Cooper et al., 2010).

Nevertheless, a significant limitation to use of BODIPY is its strong background green fluorescence even in the absence of cells, a feature repeatedly flagged as a serious impediment to microscopy studies (Cooper et al., 2010; De la Hoz Siegler et al., 2012; Govender et al., 2012). This behavior is due to the small Stokes shift between the excitation and emission maxima. In addition, the emission spectrum of BODIPY overlaps with that of green fluorescent protein (GFP), a commonly used protein fluorescent marker (Wiedenmann et al., 2009), thereby limiting its use in molecular studies. Finally, significant challenges exist in obtaining the BODIPY fluorophore. Early synthetic methods were low-yielding and led to complex mixtures (Treibs and Kreuzer, 1968) and despite subsequent advances, synthetic yields remain typically low (∼10-30%). Functionalizing the BODIPY core for specific applications is also complex: modifying the α-, β-pyrrolic, and meso-positions can be challenging and does not invariably yield the desired outcome (Boens et al., 2019). These difficulties, combined with the high cost of commercial BODIPY dyes, remain major drawbacks. Alternatives to BODIPY in algal research are hence of keen interest. Recent reports identify several new classes of fluorescent lipid-specific stains, typically optimized for examining neutral lipid accumulation and LD dynamics in mammalian cell lines (Cao et al., 2025; Fam et al., 2018; Wang et al., 2022). Very few novel fluorophores have been optimized for LD imaging in green algae (Harchouni et al., 2018; Liu et al., 2022), and none have been tested in extremophilic species.

In this work, we tested and compared the effectiveness of four fluorescent stains for LD visualization in the Antarctic extremophile *Chlamydomonas priscui*. This alga is an up-and-coming model for photosynthetic adaptations to extreme conditions, and is characterized by robust growth at low temperature, high salinity, and under extreme shading (Cvetkovska et al., 2017; Dolhi et al., 2013; Hüner et al., 2022). We compared the effectiveness of the well-characterized BODIPY (Patel et al., 2019; Rumin et al., 2015) with three fluorophores (DPAS, DAF, Cou) recently reported in mammalian lipid studies (Collot et al., 2019; Maldonado-Domínguez et al., 2014; Wang et al., 2022), but not optimized for algal work. We evaluated each stain for the following parameters: 1) suitable photophysics; 2) ease of chemical synthesis; 3) LD specificity; 4) ease of visualization in algae; and 5) cellular toxicity (Figure 1). We show that all tested fluorophores can stain LDs in this green algal extremophile, but that each has unique benefits that render it suitable for specific experimental goals. We also show that each dye has specific drawbacks that must be considered during experimental design. Our work provides new tools for studying LD dynamics and TAG accumulation in cold-water extremophilic algae, and and constitutes an essential step toward establishing these organisms as productive biotechnology platforms.

**Figure 1:**
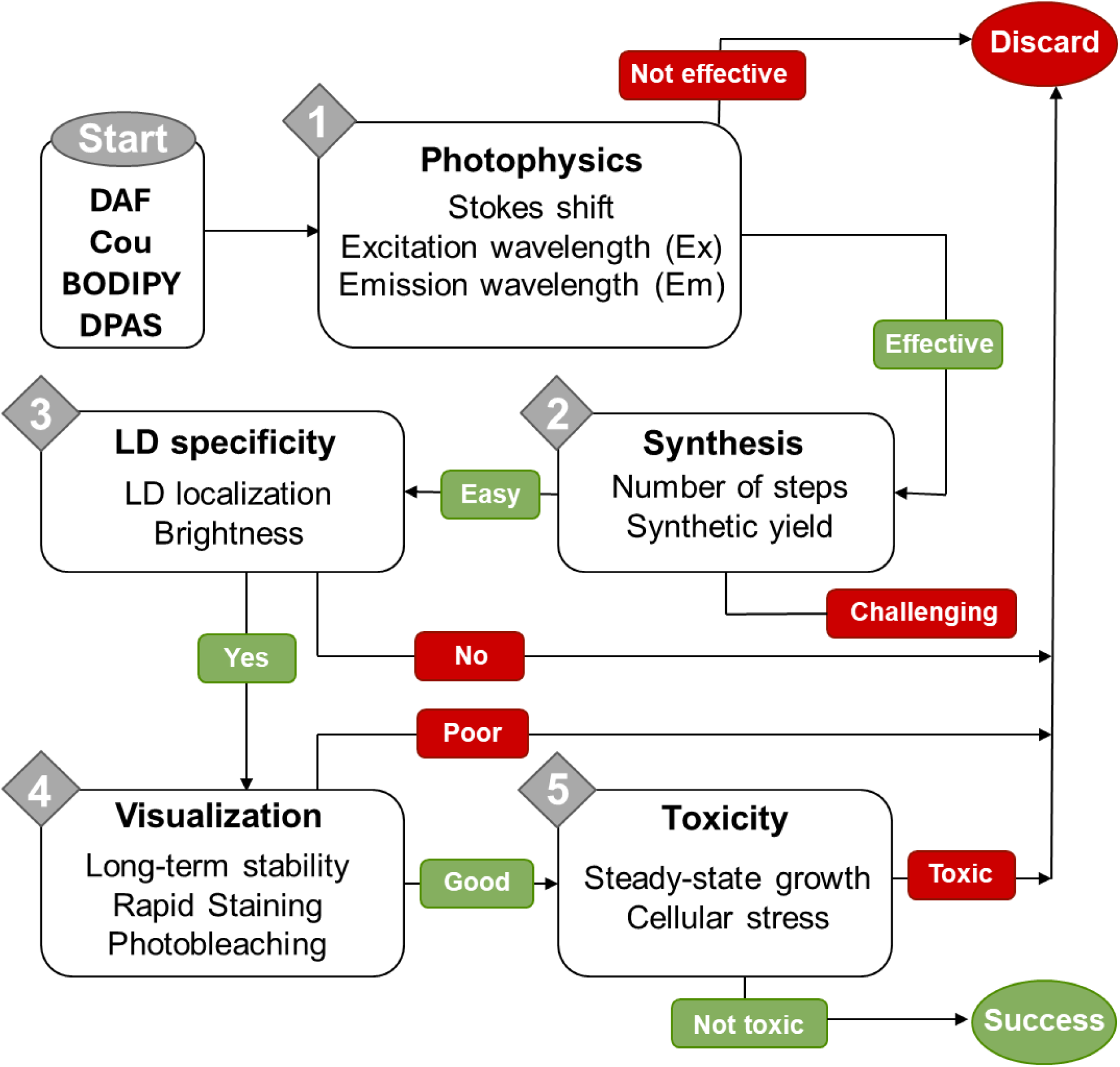
An overview of selection parameters crucial for integrating lipophilic fluorophores in typical algal workflows. All four fluorophore candidates have been used for visualizing neutral lipids in live cells, demonstrating their suitability for biological experiments, but only BODIPY has been optimized for use in green algae. The photophysical parameters and synthetic routes for each candidate fluorophore were obtained from the literature. All other parameters were tested experimentally in the present work.

## Materials & Methods

### Fluorophore synthesis

All synthesis reactions were carried out under N_2_, unless otherwise stated. Solvents employed for synthesis were distilled over P_2_O_5_ and stored under N_2_ over 4 Å molecular sieves for at least 24 h prior to use. C_6_D_6_ and CDCl_3_ (Cambridge Isotopes) were freeze/pump/thaw degassed (5×) and stored likewise. Published methods were used to prepare DPAS (2-{(*E*)-[(diphenylmethylene)hydrazono]methyl}phenol) (Wang et al., 2016), Cou (7-(diethylamino)-2-oxo-N-phenyl-2H-chromene-3-carboxamide) (Maldonado-Domínguez et al., 2014), DAF (N,N-diethylaniline furaldehyde) (Collot et al., 2019), and BODIPY 505/515 (4,4-difluoro-1,3,5,7-tetramethyl-4-bora-3a,4a-diaza-*s*-indacene) (Esnal et al., 2013; Zhang, 2017). All reagents were purchased from commercial suppliers in >99% purity. Fluorophores were stored protected from light as solids at –20 °C. Working stock solutions were prepared in DMSO, protected from light, and used immediately. The logarithm of the octanol-water partition coefficient (cLogP), used as a theoretical measure of fluorophore lipophilicity, was calculated using SwissADME (Daina et al., 2014; Daina et al., 2017).

### Algal cultivation

Axenic cultures of the Antarctic green alga *Chlamydomonas priscui* (formerly UWO241; CCMP1619) were maintained on agar slants at 4 °C. Liquid stock cultures were seeded from slants and cultivated in 250 mL Erlenmeyer flasks under the same conditions. Experimental algal cultures were grown in custom-made closed-culture bioreactors in 250 mL Pyrex tubes in temperature-regulated aquaria and aerated with sterile ambient air provided by aquarium pumps (EcoPlus EcoAir2). All algal cultures were maintained at continuous full-spectrum white light (50 μmol m^-2^ s^-1^) provided by LED lights (Ecosmart, Model # B8LT2100SGA2H04). Light intensity was measured with a quantum sensor attached to a radiometer (Model LI-189; Li-COR). In all cases, algae were cultured in Bold’s Basal Media (BBM) supplemented with 700 mM NaCl, conditions that support robust growth and closely resemble the native conditions in Lake Bonney (Antarctica) where this species was isolated (Neale and Priscu, 1995). To induce stress, algae were acclimated to BBM media supplemented with 10 mM NaCl under continuous white light (100 μmol m^-2^ s^-1^), then exposed to 24 °C for 12 hours. Such conditions were previously shown to induce cellular and physiological stress, but not cell death (Cvetkovska et al., 2022b; Possmayer et al., 2011). Unless otherwise stated, all experiments were carried out on actively growing cultures during the late logarithmic phase of growth, when the optical density at 750 nm (OD_750_) was 0.6 −1, with at least three biological replicates.

### Flow cytometry

Flow cytometry measurements (Gallios, Beckman Coultier) were collected as 20,000 events for each sample at a flow rate of 500-1000 events s^-1^. Algal cells were identified by chlorophyll autofluorescence excited at 638 nm and detected by the FL7 detector (725 ± 20 nm), eliminating cellular debris from downstream analyses. Cell size and granularity were obtained from forward-scatter (FCS) and side-scatter (SSC) measurements, respectively. Preliminary experiments were carried out to determine the optimal concentration of each lipophilic fluorophore for experimentation, defined as achieving ≥90% labelled cells within 30 minutes of adding the fluorophore (Supplementary Figure S1). In all cases, the fluorophores were dissolved in DMSO and added to algal culture in a final concentration of 1% (v/v) DMSO in BBM media. Following the addition of the fluorophores, algal cells were incubated with gentle agitation in the dark prior to measurements. The optimal working concentrations for each fluorophore was determined as follows: i) BODIPY at 2.5 μM (Ex/Em = 488/525±40 nm); ii) DPAS at 125 µM (Ex/Em = 405/550±40 nm); iii) Cou at 2.5 µM (Ex/Em = 405/450±40 nm); and iv) DAF at 50 µM (Ex = 405 nm, Em = 450±40 nm and Em = 550±40 nm for detection of fluorescence in the LDs and cytosol, respectively).

To simultaneously assess fluorophore stability and cellular toxicity, algal cultures were double-stained with both lipophilic and cell-death fluorophores. In each case, 1 mL of algal culture was incubated with each fluorophore at the concentrations indicated above for up to 24 hours in the dark with gentle agitation at 24 °C or 4 °C, as appropriate. Cell viability was assessed using SYTOX Blue (Ex/Em = 405/480 nm; Thermo Fisher Scientific, Cat. # S34857) and SYTOX Green (Ex/Em = 504/523 nm; Thermo Fisher Scientific, Cat. # S7020), which emit a fluorescent signal only when accumulating in dead cells. SYTOX was added to cell cultures at a final concentration of 1 µM, followed by incubation for 5 minutes in the dark prior to flow cytometry.

In all experiments, a control sample of cells not treated with fluorophore dye but undergoing identical incubation treatments was used to set the cutoff threshold for native fluorescence. Algal cultures treated with a single stain (either lipophilic or cell-viability dye) were used to eliminate the possibility of fluorescence overlap in double-stained experiments. Algal cells treated solely with SYTOX were used to determine cell death due to natural cell turnover in healthy cultures at 4 °C (typically ≤10%) and heat-stressed cultures at 24 °C. All data analyses were carried out using the Kaluza Analysis Software (version 2.1; Beckman Coultier). Statistical analyses were conducted in R using the rstatix package.

### Microscopy

Brightfield and fluorescence images were taken on a Zeiss AxioImager microscope fitted with an Axiocam 705 color camera using the Zen Pro Blue V. 3.2.0 software (Carl Zeiss AG). To prevent cellular motility that would interfere with image resolution, algae were embedded in low-temperature (LT) agarose prior to imaging. Thus, LT agarose (1.5% w/v) was dissolved in BBM, heated to 70 °C to dissolve the agarose, and cooled to <40 °C. Algal cultures were incubated with the lipophilic fluorophores at the concentrations indicated above for 30 minutes in the dark with gentle agitation. After incubation, stained cells were gently pelleted (3 min, 16,000 rcf), after which the supernatant was removed and replaced with cooled LT agarose. Approximately 10 μL of agarose-embedded algal culture was transferred to a glass slide, covered with a coverslip, and imaged immediately. A minimum of 10 fields of view were examined for each fluorophore.

BODIPY was excited using the 475 nm laser and detected through a 38 HE Green Fluorescent-pro filter (500-550 nm). Cou was excited with the 385 nm laser and detected through the 49 DAPI filter (420-470 nm). DPAS and DAF were excited using the 385 nm laser and detected through the 112 HE DAPI/GFP/Cy3/Cy5/Cy7 filter (412-438 nm; 501-527 nm; 582-601 nm; 662-700 nm; 770-800 nm). DPAS and DAF images were deconvoluted to remove the weak red signal arising from broad spectrum chlorophyll autofluorescence with Photoshop (v. 25.1.0; Adobe). In brief, adjustment layers were made using the channel mixer. The red channel was set to –200% (removing it entirely), while the green and blue layers remained unadjusted. The three layers were then merged using ImageJ (v 1.54i). To examine fluorophore photobleaching, videos were taken using the same preparation protocols as for the still images. Videos were recorded for 5 minutes and representative still images were recorded at key time intervals.

In the case of cellular staining with DAF, for which the emission wavelength overlaps with that of SYTOX, cell viability was measured with Trypan Blue (0.4%) (ThermoFisher Scientific, Cat. #T10282) according to the manufacturer’s instructions. In brief, equal volumes of algal culture and Trypan Blue were mixed and incubated for 5 minutes. Stained cells were mounted on a hemocytometer slide and 6 images per replicate were taken on a Zeiss AxioImager.A2 microscope (Carl Zeiss AG) fitted with a camera (model #MU1000, AmScope). Cells were manually counted in ImageJ (v 1.54i), and cell death was calculated as a percentage of blue-stained cells.

## Results and Discussion

### Four fluorophores with suitable photophysics and synthetic accessibility

To identify suitable fluorophores for LD staining in extremophilic *Chlamydomonas*, we commenced by assessing the photophysical characteristics of several dyes. First, we limited our selection to candidates that could be excited with wavelength >405 nm, to enable visualization via widely available microscopy and flow-cytometry equipment. Second, given that the photosynthetic pigment chlorophyll emits strongly in the red region of the visible spectrum (∼650-750 nm) (Donaldson, 2020), we restricted candidate fluorophores to those with emission maxima outside this range. Lastly, we targeted candidates with a Stokes shift higher than that of BODIPY 505/515 (10 nm) to address the problems arising from the small difference between excitation and emission wavelengths characteristic of many LD-specific probes (Fam et al., 2018). This final criterion is central to improving visualization quality by limiting background interference, rapid photobleaching, and poor quantification arising from fluorescence re-absorption and re-excitation.

We identified three candidates that satisfy these conditions and compared them to BODIPY (Figure 2). The first candidate, the solvatochromic fluorophore N,N-diethylaniline furaldehyde (DAF), has been used to stain LDs in human epithelial cells. DAF is excited at 390-400 nm and emits blue light (450 nm) in non-polar environments, but yellow-green (558 nm) in polar environments (Collot et al., 2019), a feature that could aid in distinguishing the effectiveness of fluorophore entry in the LDs. This dye also has a large Stokes shift (60 nm in non-polar solvents, 158 nm in polar solvents), advantageous for image acquisition. The second, 7-(diethylamino)-2-oxo-N-phenyl-2H-chromene-3-carboxamide, is an amide-functionalized, blue-emitting coumarin (Cou) derivative (Ex/Em = 422/462 nm) with a Stokes shift of 40 nm. This Coumarin fluorophore class is widely used to track enzymatic activity and tag protein biomarkers (Fan et al., 2023; Lavis and Raines, 2008) and has recently been optimized for LD visualization in mammalian cells (Xu et al., 2019; Yoshihara et al., 2020). Finally, 2-*E*-diphenylmethylene-hydrazonomethyl-phenol (DPAS), is a green-yellow emitter (Ex/Em = 390/560 nm) with a large Stokes shift (170 nm), and LD-specificity comparable to BODIPY. The latter dye was originally developed for imaging LDs in human cancer and epithelial cells (Wang et al., 2016): more recently, its use was reported in the euglenophyte alga *Euglena gracilis* (Mohsinul Reza et al., 2021). We demonstrated that all four fluorophores can penetrate the algal cell wall and accumulate in the cellular environment, as detailed below, making them viable choices for algal work.

**Figure 2:**
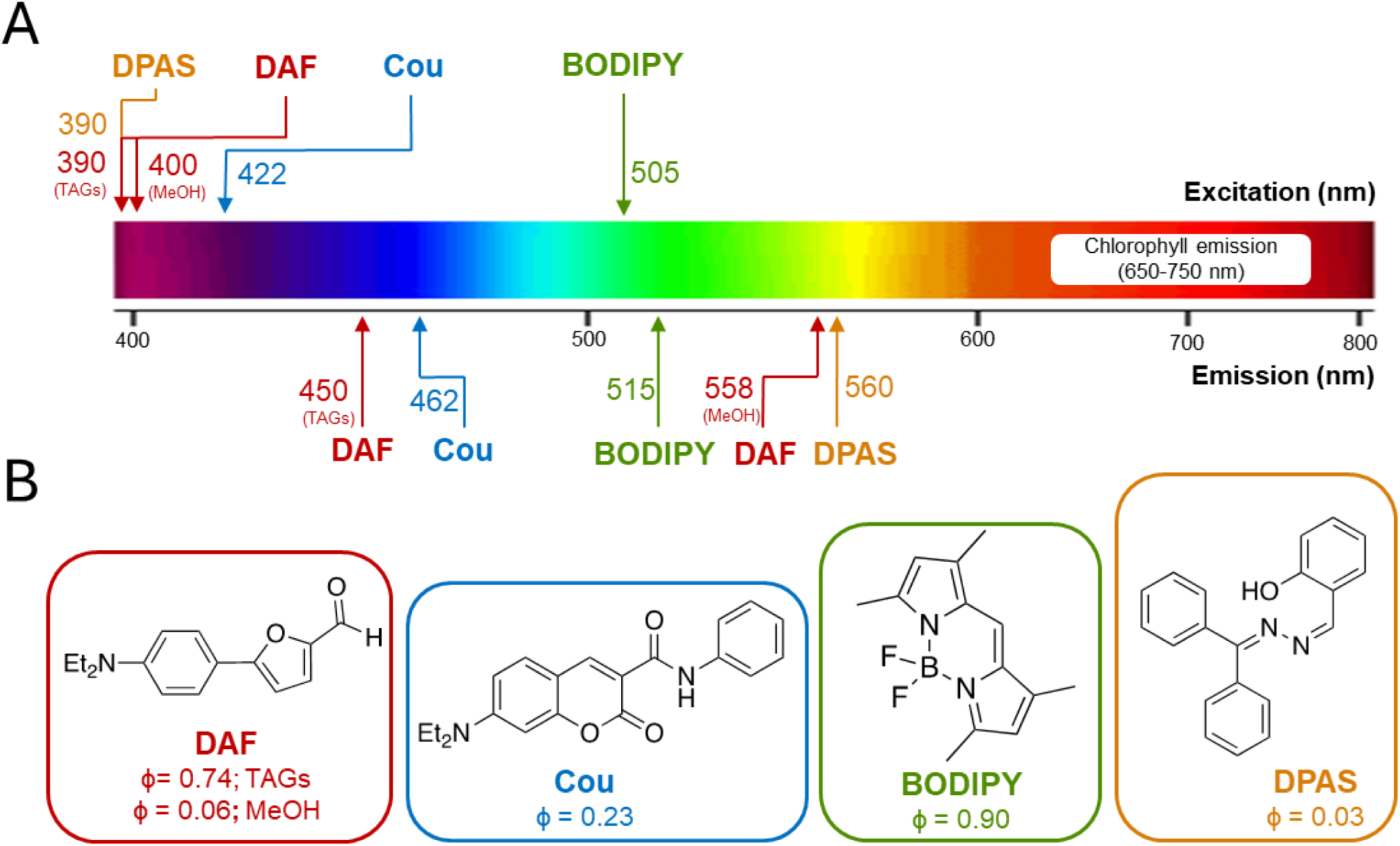
The photophysical properties of all tested fluorophores (in toluene, unless otherwise indicated). (**A**) Graphical representation of the excitation and emission bands of each fluorophore, relative to the broad chlorophyll emission band typical for photosynthetic algae. (**B**) Chemical structures and quantum yields (φ) for each fluorophore. Photophysical parameters are from published work (Collot et al., 2019; Maldonado-Domínguez et al., 2014; Rumin et al., 2015; Wang et al., 2016).

BODIPY 505/515 is commercially available, albeit costly (ca. US$300 for 10 mg; ThermoFisher). Structural modifications of the BODIPY core enables a broad array of functional derivatives used in material science and biomedical applications, including fluorescent catalysis reporters, electroluminescent films, organic light-emitting diodes, and chemosensors (Avellanal-Zaballa et al., 2022; Bumagina and Antina, 2024; Gawale et al., 2024; Li et al., 2024; Toussaint et al., 2018). These advanced applications often necessitate *de novo* synthesis of the BODIPY core, as commercially available derivatives are not readily amenable to functionalization. BODIPY is typically synthesized via a two-step reaction: the dipyrromethene core is generated *in situ* via Knorr pyrrole condensation, followed by quinone oxidation and treatment with BF_3_•OEt_2_ (Supplementary Figure 2A; Boens et al., 2019). It has been suggested that the instability of the dipyrromethane intermediate underlies the low synthetic yields (e.g., <30%; Brückner et al., 1996; Yu et al., 2003).

To circumvent these synthetic limitations, we targeted easy-to-synthesize fluorophores that have yields at least as high as BODIPY. The fluorophore DAF is synthesized over two steps (Supplementary Figure 2B), via alkylation of the precursor 4-bromoaniline, which is then appended with the furfural moiety via Suzuki coupling (Collot et al., 2019). The first step is quantitative, implicating the Suzuki coupling as the challenge in obtaining higher yields (e.g., 16% yield according to Collot et al., 2019). Nevertheless, we retained DAF as a candidate for further consideration considering the advantages offered by its solvatochromic nature. The fluorophore Cou is a functionalized derivative of coumarin, prepared from a commercial precursor via amide coupling (Supplementary Figure 2C) in a typically a high-yield reaction (e.g., 74% reported in Maldonado-Domínguez et al., 2014). While the fluorophore core for Cou must be purchased commercially, its low price (e.g., US$5/g; CombiBlocks) is suitable for widespread use. Finally, DPAS is similarly accessible in high yields (e.g., 93% reported in Wang et al., 2016), albeit in a two-step reaction from a benzophenone parent via two successive imine condensation (Supplementary Figure 2D; Wang et al., 2016).

### Optimizing LD visualization in *C. priscui*

Successful LD visualization via fluorescent dyes depends on the quantum yield and the lipophilicity of the fluorophore (Grimm and Lavis, 2022). The quantum yield (φ) is a measure of the efficiency with which a fluorophore emits absorbed photons as fluorescence. Thus, this parameter dictates the intensity of the observed signal, and the concentration of dye needed to achieve effective visualization. Lipophilicity, or the affinity of the dye for a non-polar environment, determines the specificity of the fluorophore for the algal LDs relative to polar cellular sites.

The high quantum yield of BODIPY and its derivatives (φ ∼ 0.9; Figure 2) (Loudet and Burgess, 2007) have made them attractive choices in many applications. Very low concentrations of this fluorophore (1-2.5 μM) were successfully employed to rapidly label neutral lipids in *C. priscui* cells (Supplementary Figure 1); however, while the BODIPY fluorophore localized to the LDs (cLogP = 1.57), a diffuse, non-specific cytoplasmic fluorescence (Figure 3) was also observed. Background fluorescence results in a weak correlation between BODIPY fluorescence and lipid content, a challenge that has been explicitly identified as a barrier to use of this fluorophore for quantitative lipid studies in model green algae and diatoms (De la Hoz Siegler et al., 2012; Harchouni et al., 2018). Our results indicate a similar obstacle to the use of BODIPY with the extremophile *C. priscui*.

**Figure 3:**
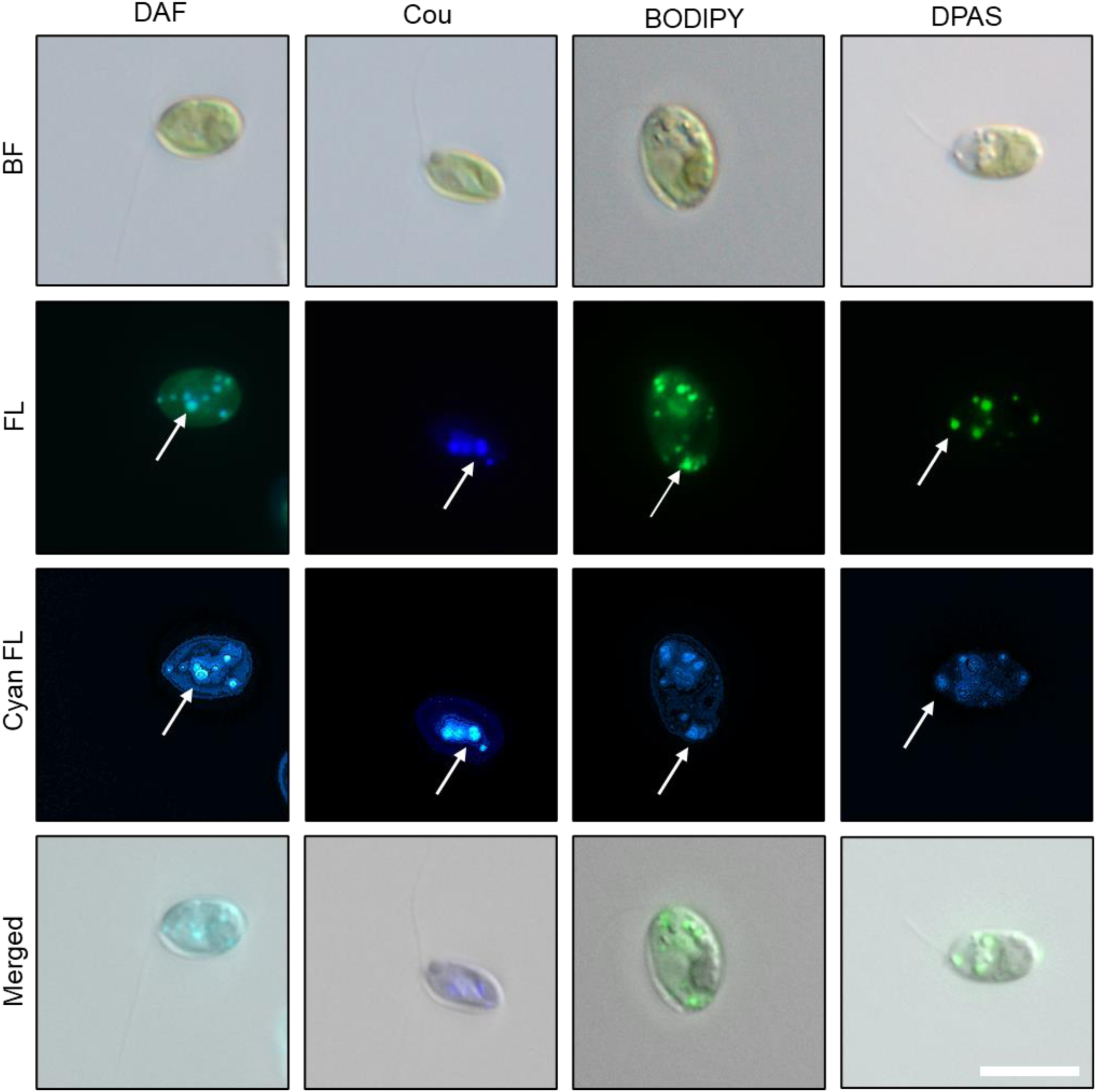
LD staining in *C. priscui*, with fluorophore localization highlighted with arrows. **BF**: Brightfield images of *C. priscui* cells grown under conditions most similar to their native environment. **FL**: True-color fluorescence microscope images of the same cells stained with DAF, Cou, BODIPY, and DPAS, showing localization of multiple LDs. **Cyan FL**: False-color (but otherwise unaltered) fluorescent microscope images, depicted for direct comparison of brightness of each dye. **Merged**: BF- and FL-merged images. In all cases, representative images are shown. Scale bar = 10 μm.

A comparison of BODIPY with DAF, Cou, and DPAS reveals several advantages of the latter three stains. Optimizing the concentration of the solvatochromic DAF presents a unique challenge, as the quantum yield of this fluorophore is higher in non-polar than polar solvents (φ = 0.74 and 0.04, respectively). In consequence (and likely related to the high lipophilicity of DAF; cLogP = 2.91), much lower concentrations were necessary to achieve comparable fluorescence in the non-polar LDs relative to the polar cytoplasm (Supplementary Figure 1, Figure 3). The DAF concentration required to stain the algal LDs (50 μM) is higher than what is required to label these organelles in HeLa cells (2 μM), pointing to less efficient entry into algal compared to mammalian cells. Despite these challenges, the solvatochromism of this dye could open unexplored opportunities in algal research. For instance, DAF was used to achieve polarity mapping in the cytoplasm and LDs in HeLa cells, and was able to report on variations in polarity between the LD core and surface (Collot et al., 2019). Detailed mapping of LD membrane composition and dynamics has not yet been accomplished in green algae, and DAF could open the door to such work.

The coumarin derivative Cou, notwithstanding its lower quantum yield (φ = 0.23; Figure 2) (Sun et al., 2018), enabled labelling of LDs in *C. priscuii* at a concentration identical to that for BODIPY (2.5 μM; Supplementary Figure 1). Reduced non-specific fluorescence was evident when compared to BODIPY (Figure 3), presumably owing to the higher lipophilicity (cLogP = 3.39) of DPAS. While the Cou fluorophore has not been used in algae, similar concentrations of coumarin derivatives have been successfully used to label LDs in mammalian cells (Xu et al., 2019; Yoshihara et al., 2020), suggesting that the algal cell wall does not function as a barrier to dye entry.

DPAS exhibits the lowest quantum yield of all tested fluorophores (φ = 0.03; Figure 2) (Wang et al., 2016), and consequently required the highest dye concentration to achieve effective LD labelling (125 μM; Supplementary Figure 1). This concentration is significantly higher than that used in human HeLa cells and the euglenophyte *E. gracilis* (10 μM; Mohsinul Reza et al., 2021; Wang et al., 2016), perhaps indicating a different mechanisms for cellular entry due to the extremophilic biology of *C. priscui.* Nevertheless, DPAS has high lipophilicity (cLogP = 4.15) and specificity for algal LDs, with minimal cytoplasmic background fluorescence compared to BODIPY (Figure 3).

### Long- and short-term staining reveals the unique characteristics of the four fluorophores

Next, we examined the stability of the fluorescent signal of the four fluorophores in aqueous media over 24 hours and correlated the effects of long-term exposure on algal viability. Staining of algal LDs with fluorescent dyes is typically a fast process (complete within 1-40 minutes) that causes no loss of viability in many species (Brennan et al., 2012; Cooper et al., 2010; Govender et al., 2012; Harchouni et al., 2018; Südfeld et al., 2021; Velmurugan et al., 2013; Xu et al., 2013) but such short-term staining does not support tracking of real-time LD dynamics in time-course experiments. To examine the long-term stability and toxicity of the dyes in algal media, we conducted simultaneous measurements of fluorescence and cell viability over a period of 24 hours. We used *C. priscui* cultures in the late-exponential phase of growth (∼9 x 10^6^ cells/mL; OD_750_ ∼1) previously shown to support high TAG accumulation in *Chlamydomonas* (Russo et al., 2017; Siaut et al., 2011). As the Antarctic *C. priscui* thrives at 4 °C (Cvetkovska et al., 2022), we chose to carry out long-term incubation experiments at this temperature. The role of temperature in fluorophore uptake is not well understood. Higher temperatures marginally decrease the uptake of BODIPY in algae (25–45 °C; Brennan et al., 2012), but no reports to date probe the effectiveness of this or other dyes in the cold. Even studies on polar *Chlamydomonas* typically employ short-term (1-5 minutes) incubation at room temperatures, and do not report uptake efficiencies at lower temperatures (Kim et al., 2016; Mou et al., 2012).

BODIPY exhibits excellent long-term stability in aqueous media in the presence of low intensity light suitable for algal cultivation and exposure to this dye has no effect on cell viability over 24 hours (Figure 4A, 4B). Thus, this stain can be successfully used for LD tracking in time-course experiments. DAF and Cou are stable in aqueous media for up to 6 hours but exhibit a small (but significant) decrease in signal intensity after 24 hours (98% and 91%, respectively; Figure 4A). Unlike Cou, which induced only a minor increase in algal mortality after 24 hours (∼8% cell death after 24 hours), DAF appears to be highly toxic to *C. priscui* cells. We observed a significantly higher proportion of dead cells even after a short incubation (∼35% cell death after 1 hour) with cellular mortality increasing substantially over the experimental period (∼50% cell death at 3 hours, and up to ∼75% after 24 hours) (Figure 4B). Several potential modes of cellular toxicity are plausible for DAF. For instance, the terminal aldehyde group readily forms Schiff bases with primary amines, such as the ε-amino group of lysine, which may disrupt protein function (LoPachin and Gavin, 2014). The conjugated π-system may also undergo light-induced redox cycling (Farmer and Davoine, 2007), generating reactive oxygen species (ROS) and leading to oxidative damage and cellular death in algae (Pérez-Pérez et al., 2012; Ugya et al., 2020; Wakao and Niyogi, 2021). In addition, the planar aromatic structure of the DAF molecule resembles that of known Photosystem II (PSII) inhibitors (e.g., phenolic herbicides). Such compounds bind at the plastoquinone QB site, blocking the photosynthetic electron flow, disrupting photosynthesis, and further enhancing ROS generation (Takahashi et al., 2010; Zharmukhamedov and Allakhverdiev, 2021). Indeed, structurally similar aldehydes, such as furfural, are known to cause oxidative stress and growth inhibition in the green algae *Scenedesmus quadricaudata* (Bringmann and Kühn, 1980) and *Chlorella vulgaris* (Kriechbaum et al., 2024). In summary, while DAF may be an excellent choice for short-term experiments, the cellular toxicity inherent to its molecular structure is a limitation, particularly for exposure beyond 1 hour.

**Figure 4:**
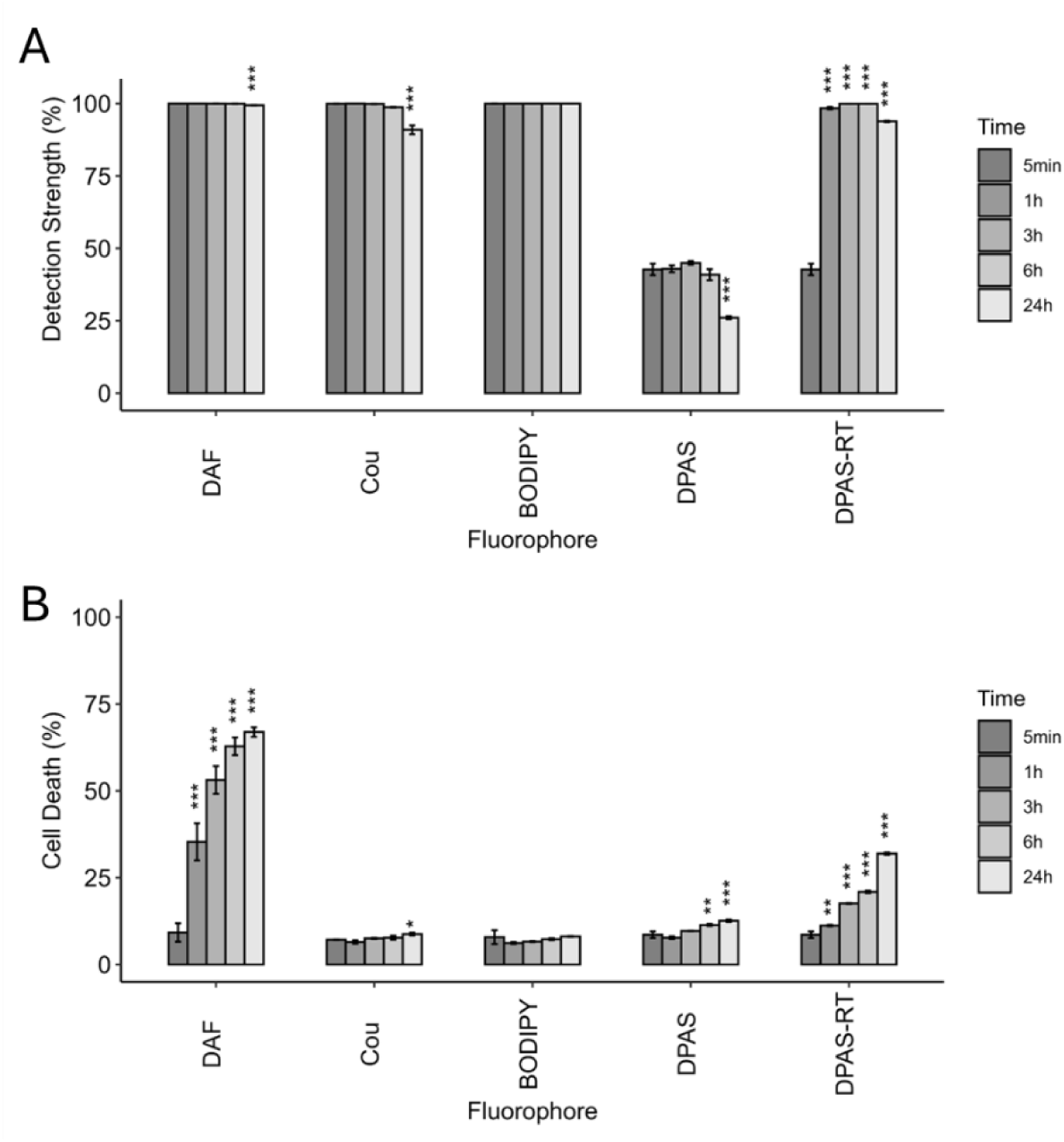
Fluorophore stability (**A**) and cellular toxicity (**B**) during long-term incubation in algal media. The fluorescence detection strength is the percentage of fluorescent cells at the specified time compared to that at the initial time-point (5 minutes). In all cases, cells were incubated with fluorophores under a gentle agitation at 4 °C, except DPAS-RT, which was incubated at room temperature. Data is average of three replicates; error bars represent the standard deviation. Statistical significance was determined by ANOVA followed by Dunnett’s post-hoc test (*** = p<0.001, ** = p<0.01, * = p<0.05)

Uniquely for DPAS, the fluorescence signal intensity and long-term toxicity proved dependent on incubation time and temperature. While DAF, Cou, and BODIPY appear to be efficiently taken up by the algal cells at 4 °C, we observed much weaker fluorescence when the cells were incubated with DPAS at this temperature (∼43% after 1 hour; Figure 4A), suggesting that uptake of this dye is inhibited in the cold. Incubating the algal cultures at 24 °C improved DPAS uptake (∼99% after 1 hour), but incubation of at least 30 minutes (not shown) was required for a strong fluorescence signal, compared to <5 minutes for the other fluorophores (Figure 4A). The temperature dependence of DPAS uptake and signal intensity may be related to its fluorescence mechanism based on aggregation-induced emission (AIE). AIE-fluorogens, such as DPAS, are non-fluorescent in solution but emit strongly as nanoaggregates (Luo et al., 2001). Monomers (or small oligomers) must dissociate from these aggregates to enter the cellular interior, a kinetically controlled process slowed by reduced thermal motion at low temperatures (Hong et al., 2011; Mei et al., 2015). This, in turn, may reduce the pool of membrane-permeable disassociated DPAS molecules and greatly decrease the overall signal intensity at low temperatures.

Incubation with DPAS also leads to small but significant decreases in cell viability after 24 hours at 4 °C, an effect exacerbated by incubation at 24 °C (∼12% at 4 °C; ∼32% at 24 °C; Figure 4B). DPAS contains multiple molecular features commonly associated with cytotoxicity. Among these are conjugated aromatic rings that can promote photosensitized ROS generation (Farmer and Davoine, 2007), a hydrazone moiety that can engage in metal-ion chelation (Sharma et al., 2020), and a planar structure that mimics PSII inhibitors (Takahashi et al., 2010). The greater bulk of DPAS may reduce its cellular toxicity relative to the more compact DAF: for example, by limiting its effective fit into the PSII binding sites. Long-term incubation of the temperate alga *E. gracilis* with high DPAS concentrations did not lead to cellular toxicity or retardation of cellular growth (Mohsinul Reza et al., 2021), indicating that the toxicity of DPAS is amplified in the polar *C. priscui*. In considering the temperature-dependent toxicity, it should be recognized that polar extremophiles are unable to survive even moderate temperatures, and *C. priscui* exhibits a heat-shock response and cell death at 24 °C (Cvetkovska et al., 2022; Possmayer et al., 2011). Increased membrane permeability and ensuing DPAS accumulation at 24 °C, combined with the effects of cellular stress on the physiology of this Antarctic alga (Cvetkovska et al., 2022a; Cvetkovska et al., 2022b) would plausibly exacerbate the toxicity of this fluorophore at the higher temperature. These results underline the importance of considering the role of incubation temperature when using fluorescent dyes to examine lipid accumulation in cold-water species.

While the photostability of BODIPY is improved relative to early-generation lipid stains, as noted above, its loss of fluorescence under high-intensity excitation light typically used in microscopy studies remains a concern (Patel et al., 2019; Rumin et al., 2015). Photobleaching (that is, the irreversible loss of fluorescence due to photochemical alteration of the fluorophore molecule) can severely impede accurate quantitation of lipid content and limit the effectiveness of microscopic observations. To probe this point, we compared the light-sensitivity and photobleaching rates of DAF, Cou, and DPAS to that of BODIPY in fluorophore-treated *C. priscui* cells. False-color images can enable direct comparison of photobleaching rates, unimpeded by the different emission spectra of the four fluorophores (Figure 2, Figure 3).

While all four fluorophores exhibited sensitivity to microscope excitation light (Figure 5; Supplementary Figure 3), BODIPY fluorescence proved most sensitive to light-induced decreases in signal intensity. The fluorescent signal from BODIPY-stained algae was nearly lost after 20 seconds of exposure, while the signals derived from DAF, Cou, and DPAS staining remained visible even after 60 seconds of exposure. BODIPY and its derivatives are well-known triplet photosensitizers. Energy transfer from the photoexcited BODIPY triplet state to molecular oxygen causes accumulation of singlet oxygen (^1^O_2_) and irreversible bond breakage in the electron-rich BODIPY core (Pederzoli et al., 2019; Wang et al., 2023). Attenuation of this effect for the other three fluorophores is plausibly due to the presence of intrinsic redox-quenching functional groups such as phenols and aromatic amines (Ali et al., 2013) or the protective effects of AIE fluorescence in the case of DPAS (Hong et al., 2011; Mei et al., 2015). Therefore, while BODIPY may be well-suited for LD quantification via flow cytometry, the other three fluorophores are superior choices for microscopy.

**Figure 5:**
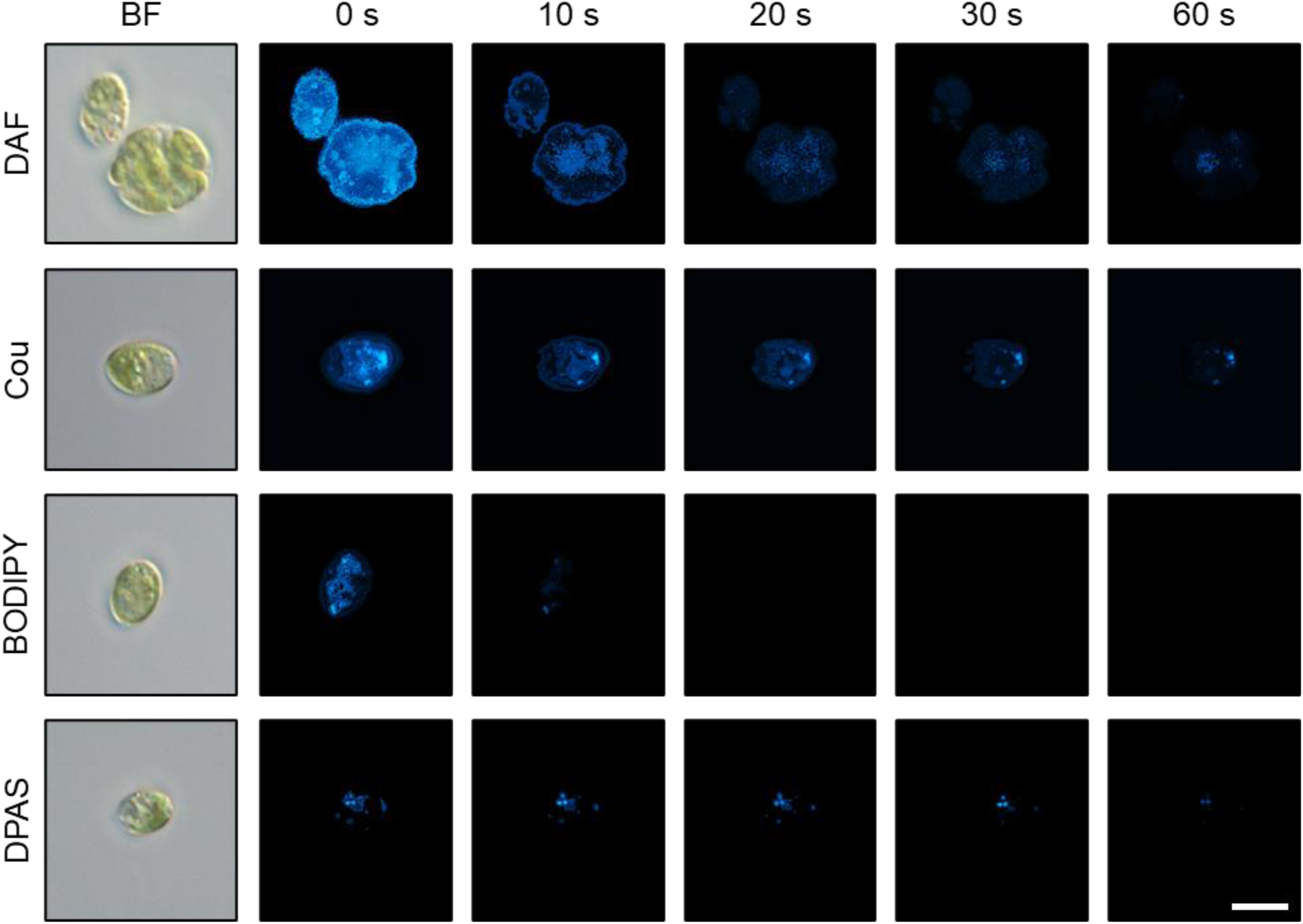
Image stills from videos demonstrating photobleaching of DAF, Cou, BODIPY, and DPAS in *C. priscui*. Time-points represent the total amount of time that the stained algal cells were exposed to microscope excitation light. Algal cells were incubated with the fluorophores for 30 minutes at room temperature and were kept in dark prior to visualization. Fluorescent images are presented in cyan false color (but are otherwise unaltered) for a direct comparison of the brightness of each dye. True color images are provided in Supplementary Figure 3. **BF**: Brightfield images of *C. priscui* cells. Representative images are shown. Scale bar = 5 μm.

### BODIPY and DAF are not suitable for studying stress-induced lipid accumulation

The synthesis of TAGs in response to stress in *Chlamydomonas* is well documented. High lipid accumulation that accompanies physiological changes has been linked to nutrient deficiencies (Davey et al., 2014; Juergens et al., 2016; Moellering and Benning, 2010; Siaut et al., 2011; Urzica et al., 2013), high light stress (Goold et al., 2016), and in response to elevated temperatures (Hemme et al., 2014; Légeret et al., 2016). The use of fluorescent dyes to study TAG turnover and LD dynamics is a common tool in such studies but reports on the effect of the dye on algal physiology or viability during stressful conditions are exceptionally rare. Indeed, fluorophores are widely presumed to be non-toxic (Cao et al., 2025; Patel et al., 2019), notwithstanding the paucity of reports assessing dye toxicity during stress in algae. In a recent exception, the observation of increased cellular death in BODIPY-stained stramenopiles during cellular stress was attributed to the exacerbating effect of BODIPY on algal stress physiology (Südfeld et al., 2021). Whether such effects are general for other fluorescent dyes is unknown, despite the critical importance of this question for the interpretation of data in algal stress studies.

We examined the toxicity of the four fluorophores in *C. priscui* cultures acclimated to optimal temperature (4 °C) and exposed to non-permissive temperature (24 °C) for 12 hours. This seemingly innocuous treatment induces heat-stress responses in this Antarctic alga at the level of photosynthesis, respiration, gene transcription and metabolite accumulation within 6 hours, although measurable cell death occurs only on exposures of >24 hours (Cvetkovska et al., 2022b; Possmayer et al., 2011). To avoid confounding effects from heat-induced mortality, we performed all experiments after 12 hours of heat stress and confirmed that algae undergoing this treatment only exhibit <5 % mortality, attributed to normal cell turnover (Control; Figure 6). In all cases, cells were treated with fluorophores for 30 minutes, a short-duration incubation that enables effective staining with all dyes but does not cause loss of viability in *C. priscui* during low-temperature growth (Figure 4B).

**Figure 6:**
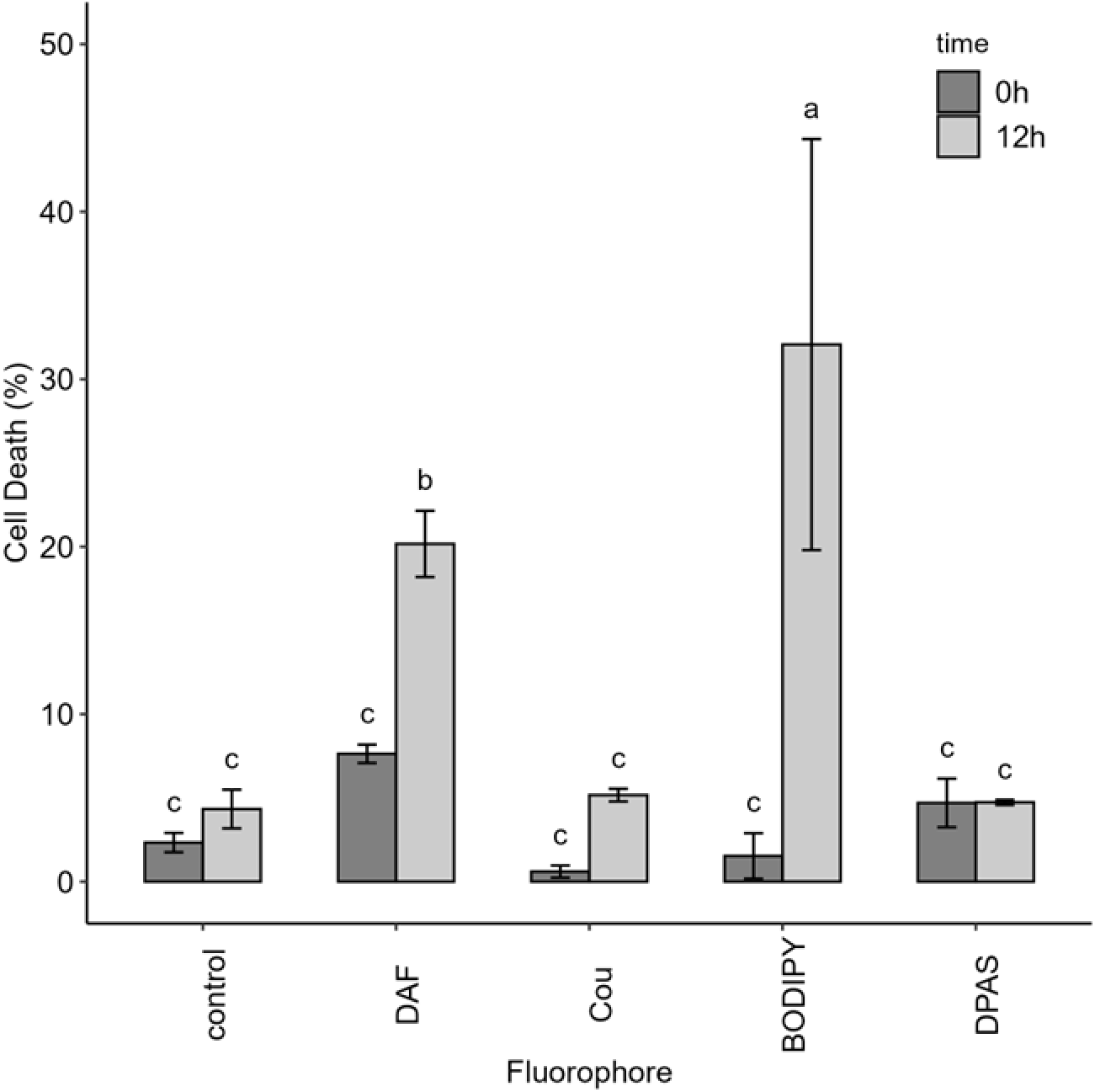
Cell death in *C. priscui* cultures at 4 °C (0 hours) exposed to short-term heat stress at 24 °C (12 hours), then stained with a lipophilic fluorophore, DAF, Cou, BODIPY, or DPAS. As a control, cultures were exposed to the same treatments but not stained. Cell death was quantified with SYTOX or Trypan Blue dead-cell indicators. Data is average of three replicates; error bars represent the standard deviation. Statistical significance was determined by ANOVA followed by Tukey’s post-hoc test, where different letters represent significant differences between treatments.

Our results demonstrate that treatment with DAF and BODIPY induces significant cellular death (∼21% and ∼31% mortality, respectively), but only in heat-stressed *C. priscui* cultures (Figure 6). In contrast, no increase in cell death was observed in heat-stressed cultures treated with Cou and DPAS (∼5% mortality in both cases). The toxicity of DAF appears to be inherent to the molecular structure of this fluorophore, as increased cellular death is observed at both low temperatures (Figure 4B) and in combination with heat stress (Figure 6). In contrast, the toxicity of BODIPY appears to be associated only with heat stress (Figure 6), as no increase in mortality is observed during 24-hour exposure during low-temperature growth (Figure 4B). The robust antioxidant system of *C. priscui*, an adaptation to life under extreme conditions (Cvetkovska et al., 2022b; Stahl-Rommel et al., 2021), may effectively counteract BODIPY-generated ROS accumulation under steady-state (low temperature, non-stressful) conditions that support optimal growth. However, temperature-sensitivity of this cold-water alga (Cvetkovska et al., 2022b; Possmayer et al., 2011) appears to increase its susceptibility to BODIPY cytotoxicity under heat stress. A correlation between loss of viability and BODIPY treatment during nutrient stress was demonstrated in the stramenopile *Nannochloropsis oceanica* (Südfeld et al., 2021), but no reports yet describe the repercussions of cellular stress on BODIPY staining in *Chlamydomonas*. Here we clearly show that the physiological condition of the cells must be taken into consideration when selecting a fluorophore for lipid detection, to prevent inaccurate assessment of cell physiology and lipid levels.

## Summary

The foregoing is the first report of LD labelling in the extremophilic green alga *C. priscui,* an up-and coming model for photosynthetic adaptation to life at the extreme (Cvetkovska et al., 2017; Hüner et al., 2022). All four tested fluorophores (DAF, Cou, BODIPY, and DPAS) localize to the algal LDs, but each presents unique advantages or limitations depending on the experimental context. BODIPY, while widely used as an excellent dye for short- and long-term staining, is highly susceptible to photobleaching, and exhibits non-specific fluorescence. This has been flagged as a serious impediment to the use of BODIPY in microscopy studies or for quantifying lipid content (Cirulis et al., 2012; Xu et al., 2013). Importantly, the negative effect of BODIPY on cell viability may also limit its applications in studies involving environmental stress. In comparison, DAF provides unique advantages arising from its solvatochromism and could be particularly useful in assessing LD entry efficiency, especially if conjugated to other molecules. However, this fluorophore must be used with caution during long-term staining or stressful conditions, as it exhibits strong cytotoxicity in *C. priscui*. The highly lipophilic Cou and DPAS provide strong LD-specific signals and minimal effects on cell viability in long-term staining experiments. However, DPAS must be used at room temperature, which limits its applicability for labeling LDs in cold-water extremophiles. These two dyes are best suited for experiments involving stress conditions, as they do not amplify endogenous cell-death response. Systematic studies confirming linear correlations between fluorescence and biochemically quantified lipid content are now under way, with the goal of establishing the power of Cou and DPAS as biochemical tools.

This work underscores the critical importance of experimental optimization in LD labelling experiments. Lipid-focused algal studies using fluorescent dyes other than BODIPY remain limited, only a few alternatives having been explored to date (Cirulis et al., 2012; Harchouni et al., 2018; Mohsinul Reza et al., 2021). We show that the physiological state of the labelled cell, and not merely optimal dye concentration and incubation time, must be considered to ensure data quality and accurate interpretation. Cold-water extremophilic algae are beginning to garner strong interest as novel targets for biotechnology applications, but very few species have been characterized in detail. Resources like the CCCryo Culture Collection of Cryophilic Algae (Leya, 2022) and advances in low-temperature cultivation of oleaginous species (Cheregi et al., 2019; Kvíderová et al., 2017; Pankratz et al., 2017; Vona et al., 2018) will undoubtedly become increasingly important as the field develops. Careful optimization of the experimental tools used to study lipids in extremophiles is essential for both fundamental understanding and practical applications. Judiciously chosen fluorophores are anticipated to offer powerful tools to probe lipid dynamics, cellular physiology, and synthetic processes in extremophilic algae.

## Acknowledgements

We acknowledgke the support from the Government of Canada’s New Frontiers in Research Fund (NFRF) [NFRFE-2022-00608], a tri-agency initiative of the Canadian Institutes of Health Research (CIHR), the Natural Sciences and Engineering Research Council (NSERC), and the Social Sciences and Humanities Research Council (SSHRC). MC, CNB and DEF are grateful for support via NSERC Discovery Grants and the Canada Foundation for Innovation (CFI). The Government of Ontario is thanked for an Ontario Graduate Scholarship to EJYB.

## Author Contributions

EJYB synthesized all fluorophores. PO performed all algal work and data analysis, including microscopy and flow cytometry. DEF and MC provided funding, supervision, data curation, and technical guidance. CNB provided technical expertise and guidance. All authors were involved in conceptual development of the work, manuscript writing and revisions, and final approval.

## Data Availability Statement

The data is available from the corresponding author upon reasonable request.

## Supporting Information

**Supplementary Figure S1**: Optimizing the concentration of (**A**) DAF, (**B**) Cou, (**C**) BODIPY, and (**D**) DPAS required to label LDs in a late-exponential culture of *C. priscui* with a high lipid content. The data shows the proportion of cells with fluorescent signals detected by flow cytometry. The optimal concentration for subsequent experiments is defined as achieving ≥90% labelled cells. Representative results are shown.

**Supplementary Figure 2: Supplementary Figure 2:** Reported routes for synthesis of BODIPY (**A**), DAF (**B**), Cou (**C**), and DPAS (**D**). In all cases, the reported synthetic yields are provided in brackets. pTsOH = p-toluenesulfonic acid; TFAA = trifluoroacetic acid; HBTU = hexafluorophosphate benzotriazole tetramethyl uranium; DMF = dimethyl formamide. In all cases, synthesis reactions have been reported previously (Collot et al., 2019; Cunha Dias de Rezende et al., 2014; Maldonado-Domínguez et al., 2014; Wang et al., 2016)

**Supplementary Figure 3:** Representative image stills from videos demonstrating photobleaching of DAF, Cou, BODIPY, and DPAS in *C. priscui*. Time-points indicate the total exposure time of the stained algal cells to microscope excitation light. Algal cells were incubated with fluorophores for 30 minutes at room temperature and kept in dark prior to visualization. BF: Brightfield images of *C. priscui* cells. All other images are true-color fluorescent images. Representative images are shown. Scale bar = 5 μm.

