## Supplementary Figure for "A Suite of Stains: Characterization of four fluorophores as complementary tools for visualizing neutral lipids in an extremophilic green alga"

### SUPPLEMENTARY FIGURES

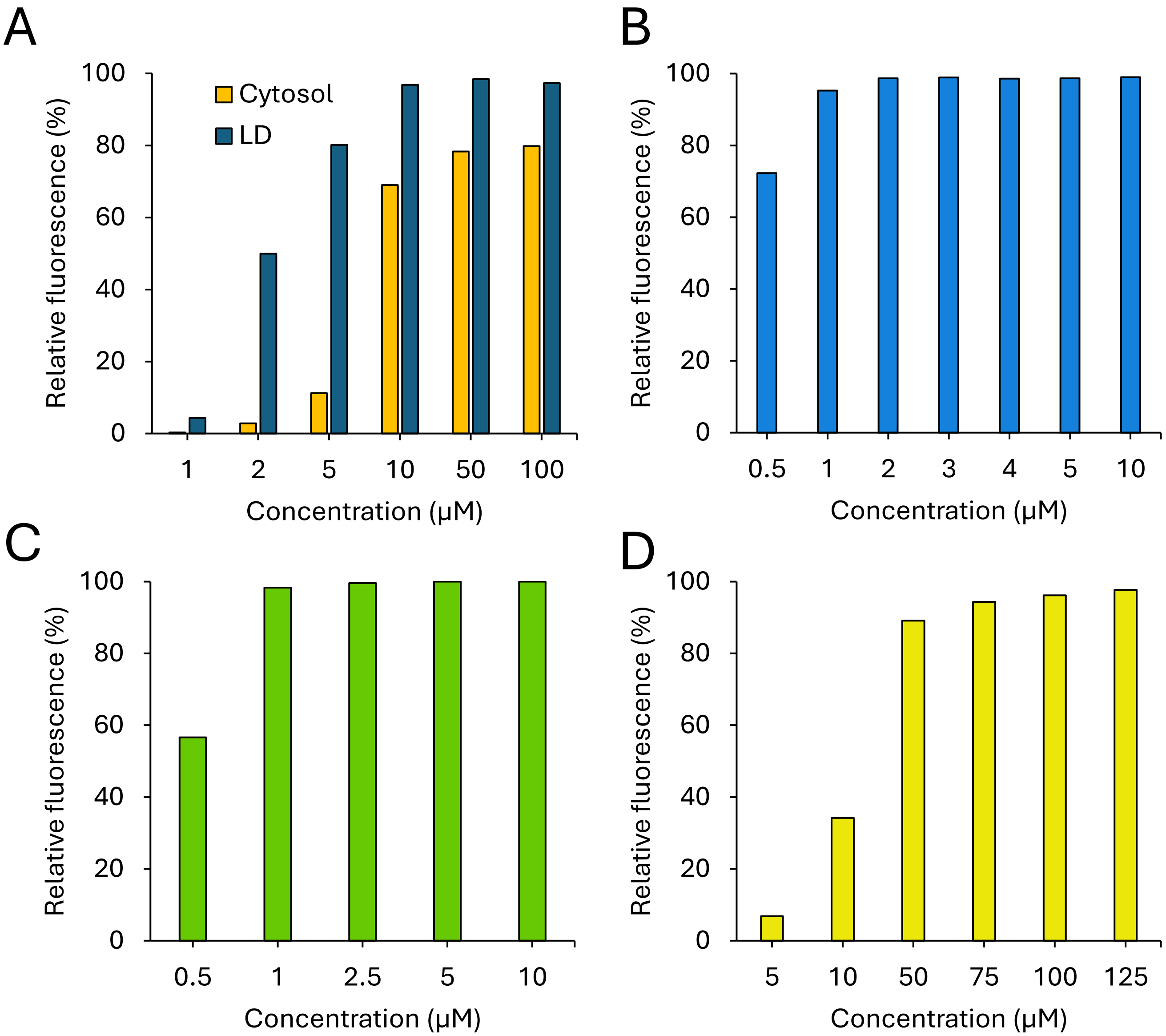


**Supplementary Figure S1:** Optimization of the concentration of (**A**) DAF, (**B**) Cou, (**C**) BODIPY, and (**D**) DPAS required to label LDs in a late-exponential culture of *C. priscui* with a high lipid content. The data shows the proportion of cells with detectable fluorescent signal detected by flow cytometry. Optimal concentration used in subsequent experiments is defined as achieving ≥ 90% labelled cells. Representative results are shown.

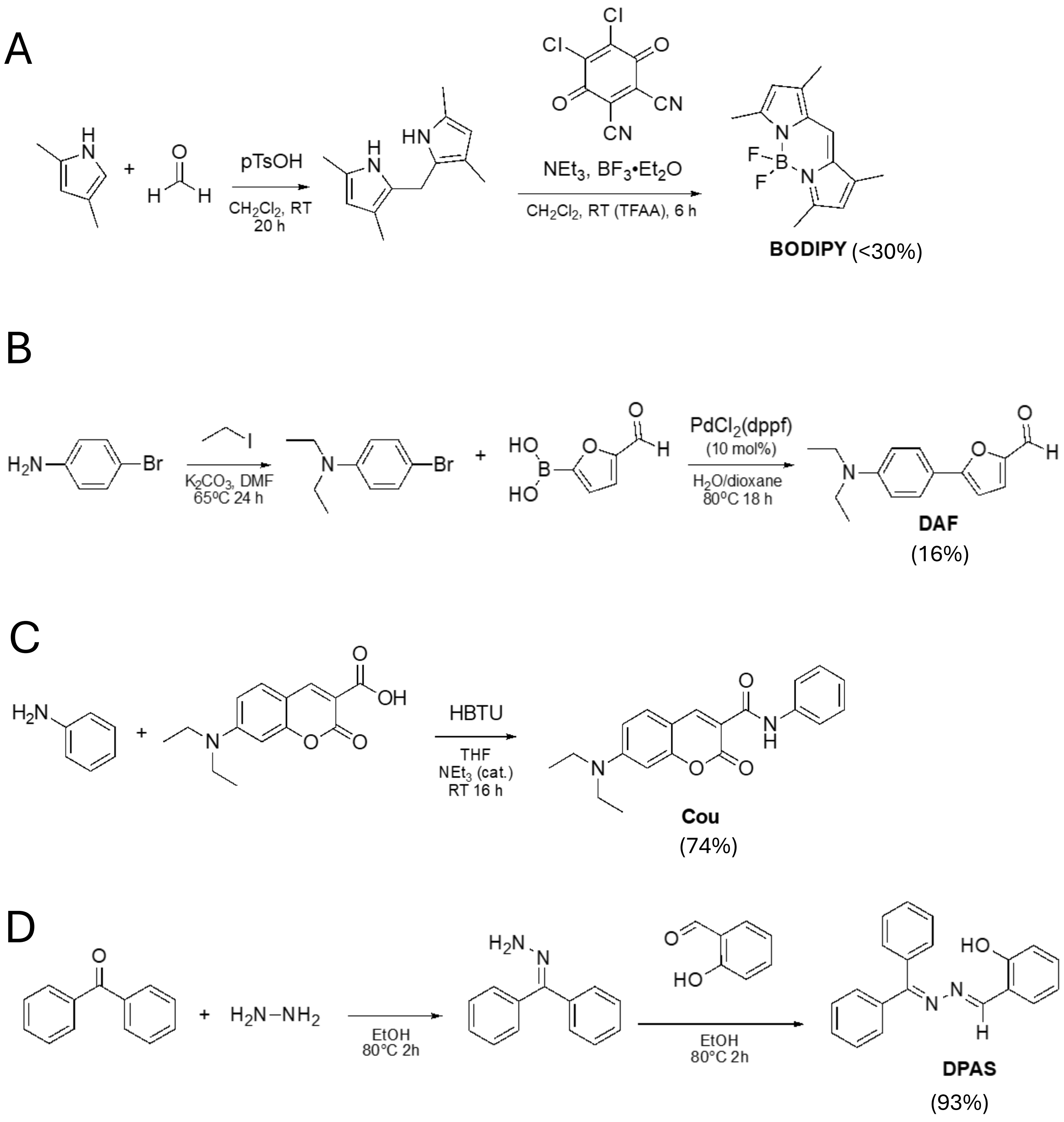


**Supplementary Figure 2**: Reported routes for synthesis of BODIPY (**A**), DAF (**B**), Cou (**C**), and DPAS (**D**). In all cases, the reported synthetic yields are provided in brackets. pTsOH = p-toluenesulfonic acid; TFAA = trifluoroacetic acid; HBTU = hexafluorophosphate benzotriazole tetramethyl uranium; DMF = dimethyl formamide. In all cases, synthesis reactions have been reported previously (Collot et al., 2019; Cunha Dias de Rezende et al., 2014; Maldonado-Domínguez et al., 2014; Wang et al., 2016)

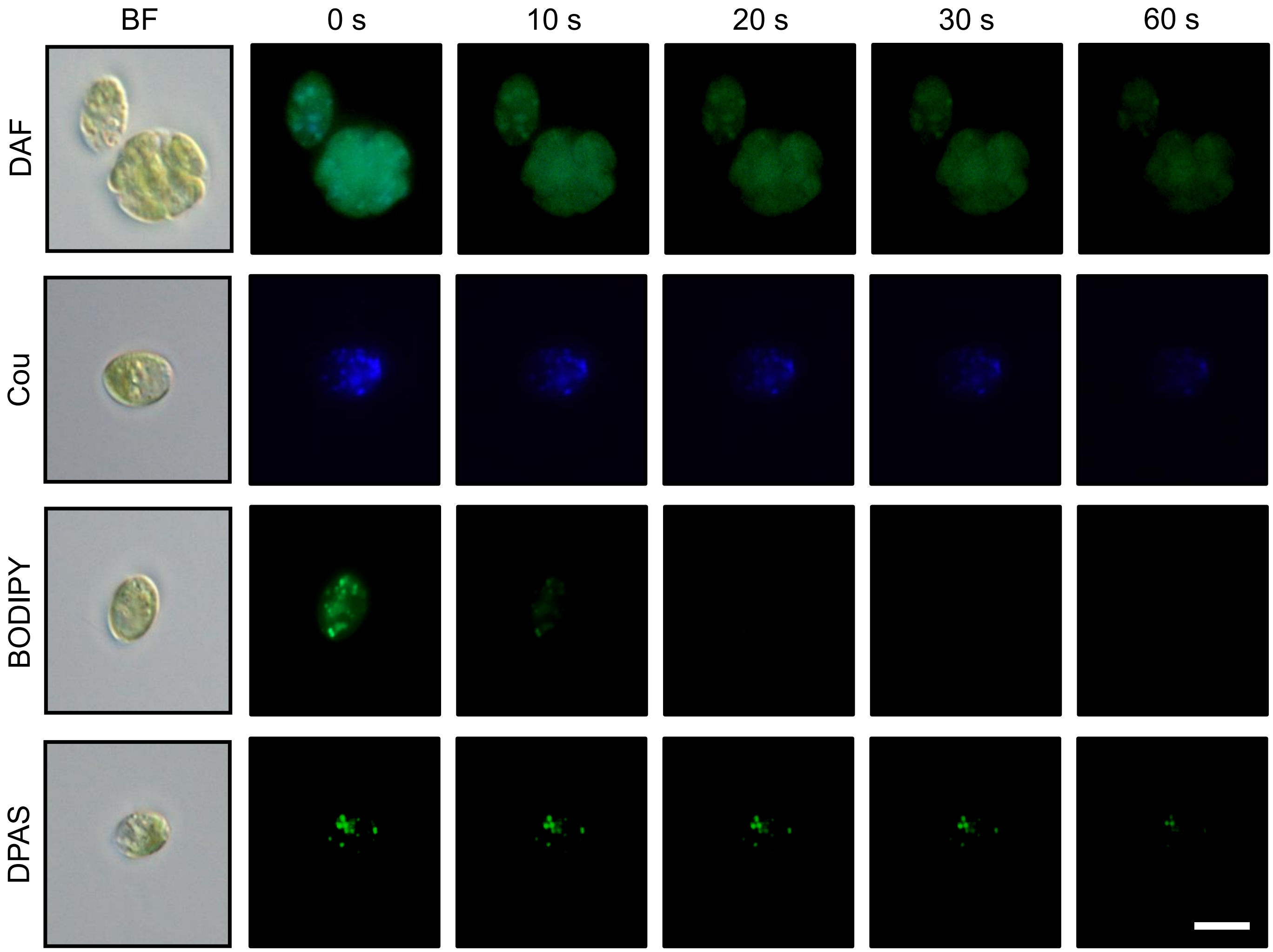


**Supplementary Figure 3**: Image stills from videos demonstrating the photobleaching of DAF, Cou, BODIPY, and DPAS in *C. priscui*. The time points represent the total amount of time that the stained algal cells were exposed to microscope excitation light. Algal cells were incubated with fluorophores for 30 minutes at room temperature and were kept in dark prior to visualization. BF: Brightfield images of *C. priscui* cells. All other images are true-color fluorescent images. Representative images are shown. Scale bar = 5 μm.
